# MaPPeRTrac: A Massively Parallel, Portable, and Reproducible Tractography Pipeline

**DOI:** 10.1101/2020.12.23.424191

**Authors:** A collaboration between the U.S. Department of Energy and TRACK-TBI, Joseph Moon, Peer-Timo Bremer, Pratik Mukherjee, Amy J. Markowitz, Eva M. Palacios, Lanya T. Cai, Alexis Rodriguez, Yukai Xiao, Geoffrey T. Manley, Ravi K. Madduri

## Abstract

Large-scale diffusion MRI tractography remains a significant challenge. Users must orchestrate a complex sequence of instructions that requires many software packages with complex dependencies and high computational cost. We developed MaPPeRTrac, a probabilistic tractography pipeline that simplifies and vastly accelerates this process on a wide range of high performance computing (HPC) environments. It fully automates the entire tractography pipeline, from management of raw MRI machine data to edge density imaging (EDI) of the structural connectome. Dependencies are containerized with *Docker* or *Singularity* and de-coupled from code to enable rapid prototyping and modification. Data artifacts are strictly organized with the *Brain Imaging Data Structure* (BIDS) to ensure that they are findable, accessible, interoperable, and reusable following FAIR principles. The pipeline takes full advantage of HPC resources using the *Parsl* parallel programming framework, resulting in the creation of connectome datasets of unprecedented size. MaPPeRTrac is publicly available and tested on commercial and scientific hardware, so that it may accelerate brain connectome research for a broader user community.

## 1. Introduction

Diffusion MRI can be digitally processed into a tensor field describing white matter fiber orientation *in-vivo* (Basser et al., 1994). By Monte Carlo sampling on this field, it is possible to estimate axon pathways between regions of the brain (Behrens et al., 2003). The resulting graph, known as a structural connectome (Sporns et al., 2005), provides a quantitative measure of brain connectivity useful to analyze both healthy brains and changes caused by neurological and psychiatric diseases (Sporns, 2013). Structural connectomes can be evaluated using a plethora of well-developed techniques based on graph theory and matrix analysis. They can be further processed into edge density imaging (EDI), which maps the number of connectome edges that pass through every white matter voxel in the brain (Owen et al., 2015, 2016; Qi and Arfanakis, 2021). Both structural connectomes and EDI show great promise towards the timely evaluation of disorders of white matter connectivity such as traumatic brain injury (TBI) (Raji et al., 2020; Reber et al., 2021).

Recent advances in diffusion MRI have enabled the creation of vast high-resolution datasets (Palacios et al., 2020). However, connectome research has been limited by the throughput of neuroimaging software. Existing tools tend to require complicated prerequisite tools that are difficult to install and run or that do not support parallel and efficient execution. Generating connectomes for even a handful of patients can take days on a high-end personal computer and requires the dedicated attention of an operator skilled in neuroimaging and computer science (Madhyastha et al., 2017). As a result, dealing with cohorts of hundreds or thousands of patients is virtually impossible for most research groups. Addressing this challenge requires a neuroimaging pipeline that can be easily deployed, has been designed to use high performance computing (HPC) seamlessly, and can leverage existing standards to ensure reproducibility. And although other pipelines can generate structural connectomes, at time of publication this pipeline will be the only publicly available software that can generate EDI at scale.

### 1.1. Related Work

There exist several neuroimaging pipelines for diffusion MRI, but only a few take advantage of HPC capabilities. This is especially true at the largest scale, where computation is coupled to sophisticated resource allocation algorithms and distributed across multiple different clusters. The LONI Processing Environment provides an interactive neuroimaging workspace with fully modular components (Rex et al., 2003). However, it has several limitations: HPC execution is only possible with the Grid Scheduler, heterogeneous resources such as GPUs are poorly supported, and software dependencies must be installed manually with elevated-user privileges.

Schirner et al. (2015) propose a multi-modal pipeline that can generate connectomes on HPC clusters. Though closely related, it does not meet all of our requirements. The pipeline’s imperative software architecture makes data and scripts brittle and difficult to modify. There is no support for GPU acceleration or low-level parallelism, with the result that processing even a single patient exceeds time limitations on certain HPC clusters. And similar to the LONI pipeline, it demands root privileges to install software libraries. There exist functional MRI (fMRI) pipelines that *do* meet many of our computational requirements, such as *TractoFlow* (Theaud et al., 2020) and *NDMG* (Kiar et al., 2017). However, none of them can be configured to run probabilistic tractography on structural connectomes. This is particularly limiting as new research suggests that EDI generated using structural connectomes offer novel insights distinct from fMRI connectomes (Reber et al., 2021). Furthermore, the tight software coupling of fMRI pipelines makes them impractical to re-write to our requirements.

It is important to distinguish neuroimaging pipelines from tractography software tools, which are generally run on personal computers. There exists dozens, if not hundreds, of the latter, each with various software configurations and performance characteristics (Côté et al., 2013; Maximov et al., 2019; Cui et al., 2013). However, these tools are nearly all aimed towards processing individual patients and cannot take advantage of HPC parallelization. And similar to non-modular pipelines, tractography software tools often suffer from complex dependencies that present obstacles to non-expert users.

### 1.2. Requirements

We propose a novel probabilistic tractography pipeline for structural connectomes. When designing this pipeline, several competing needs have been considered.

#### 1.2.1. Need for high performance

Analysis methods should be designed and implemented using appropriate software constructs to take advantage of HPC resources. This is especially the case for pipeline instructions that have a memory footprint larger than what personal computers can handle. The Department of Energy is home to some of the world’s fastest supercomputers, opening a unique opportunity for large-scale tractography (Top500, 2020). Parallelization and high performance go hand in hand, so understanding the inherent parallelization in the analysis will lead to vastly increased efficiency. One must pay close attention to potential parallelism at every step in the tractography pipeline, so that it may reduce the steep computational cost.

#### 1.2.2. Need for reproducibility

Regardless of performance, any tractography pipeline must produce results that are consistent across multiple execution environments and accurately represent the intentions of the analysts. For probabilistic tractography, this can be divided into processes that are deterministic and processes that are stochastic in nature. Deterministic algorithms, such as brain extraction and segmentation, should produce identical outputs given same inputs on all computing environments. Any stochastic process - in this case, the tractography itself - must produce results that are roughly similar given identical inputs, with values converging as the number of samples increases. Reproducibility is particularly important for connectome analysis, since it remains difficult to validate the findings of graph theoretical methods merely with neurological observation (Roine et al., 2019).

#### 1.2.3. Need for portability

We define portability as the ability to run on any available computational resources and environments. Portability is key for reusability as well as wider adoption and deployment. All of the high performance optimizations should not be limited to HPC platforms, but potentially run on most Linux environments (even if they do not have the horsepower to run tractography efficiently). It is also important the input and output data be easily shared between users. As far as possible, data should be organized in standard, machine-readable structures and use consistent file formats. Similarly, community standards and best practices should be used for metadata and naming conventions, so that individual files can be more easily identified and manipulated. Lastly, the design and implementation of the pipeline should follow best practices of object oriented programming techniques and enable customization by configuration files so the pipeline can easily be extended and modified to suit different users and sites.

### 1.3. Summary of MaPPeRTrac contributions

Our objective is to perform tractography analysis - from raw signal processing to generating the final connectome and EDI - while taking full advantage of HPC resources. Code and dependent libraries ought to be portable across computing systems, including those where root privileges are not available. Neuroimaging parameters and resource allocation should be configurable and consistent across different software components. Deterministic algorithms must be reproducible across systems. And most significantly, the pipeline should be as performant as possible in a parallel computing environment. To this end, we have developed MaPPeRTrac, a **M**assively **P**arallel, **P**ortable, and **R**eproducible **Trac**tography pipeline that is capable of:

1. Leveraging parallelization: Take advantage of the inherent parallelism in neuroimaging analysis, not only across participants but also within individual analyses;
2. Portability across computing infrastructures: Transparently execute neuroimaging analysis on computing infrastructures at different institutions, with minimal changes to the pipeline itself.
3. Reproducing deterministic results: Ensure that results at different computing sites are exactly equivalent (or closely comparable, for stochastic procedures such as tractography).
4. Configuring all neuroimaging parameters: Allow easy access to the various parameters used by neuroimaging components in the pipeline; and
5. Being FAIR: Ensure that datasets, software, and other digital objects are **F**indable, **A**ccessible, **I**nteroperable, and **R**eusable according to standard FAIR guidelines (Wilkinson et al., 2016). This facilitates long-term use in research, promotes knowledge integration, and increases reusability of existing data.

## 2. Background and Materials

### 2.1. Original Tractography Scripts

Our pipeline emerged from an existing collection of neuroimaging scripts by (Owen et al., 2015, 2016). These scripts produce two outputs: 1) the grey matter connectivity adjacency matrix typical of a structural connectome and the EDI associated with the structural connectome. These are generated through the following steps:

1. Manually convert DICOM scanner images into NIfTI format using *dcm2niix*
2. Read NIfTI-formatted diffusion MRI data. Correct for motion and scan artifacts, remove non-brain tissue, and calculate diffusion anisotropy.
3. Estimate diffusion parameters and fiber directions in each voxel
4. Parcellate the cortical and subcortical gray matter into 82 regions based on the Desikan–Killiany atlas (Desikan et al., 2006)
5. Compute white matter fiber streamlines to generate a structural connectome using probabilistic tractography. Collate tractography into an EDI map (e.g. Figure 1).

**Figure 1:**
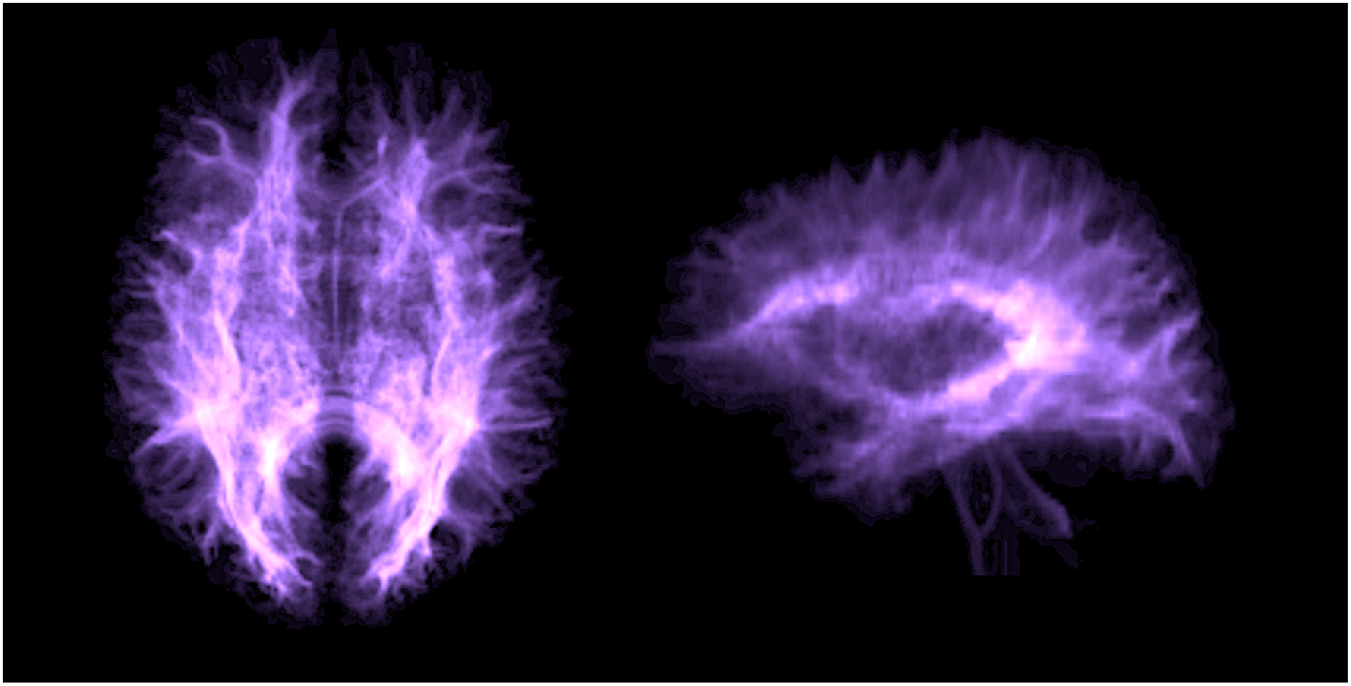
An edge density image (EDI) generated by MaPPeRTrac

**Figure 2:**
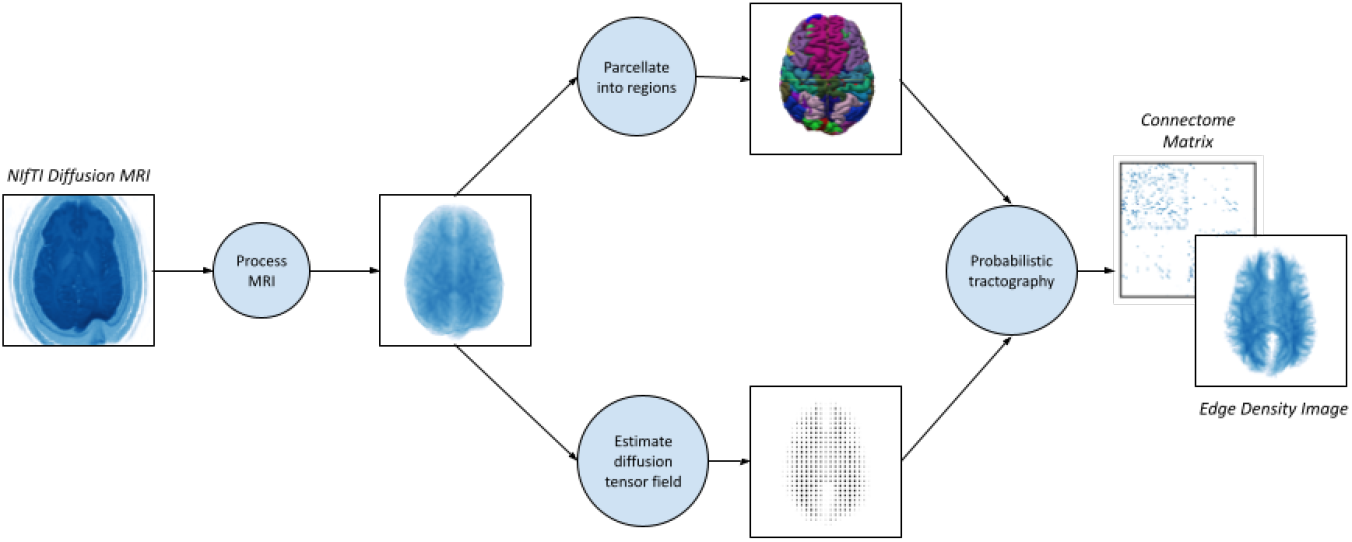
Data artifacts from the tractography workflow (both original scripts and MaP-PeRTrac)

**Figure 3:**
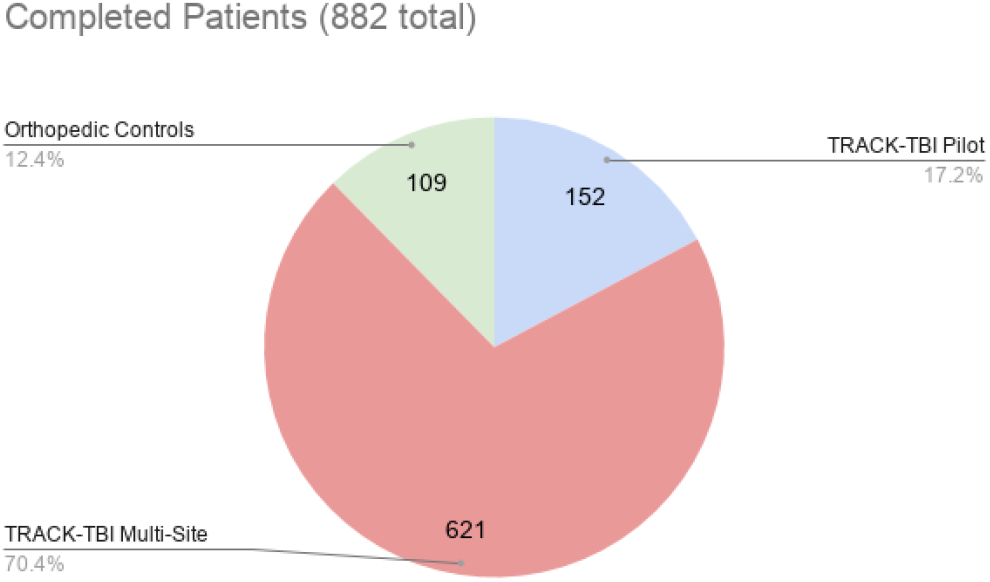
Breakdown of patient population

**Figure 4:**
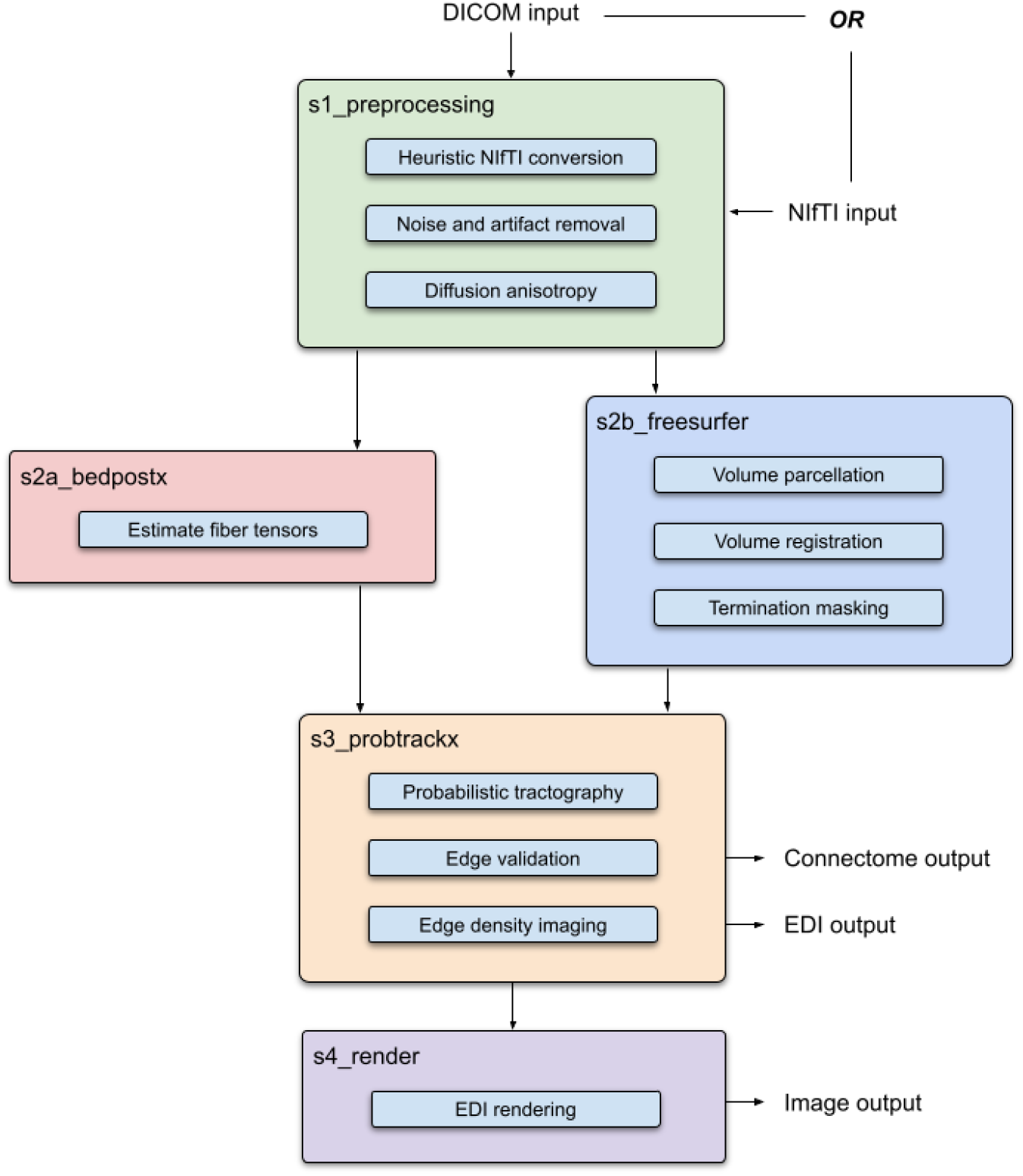
Overview of MaPPeRTrac architecture

**Figure 5:**
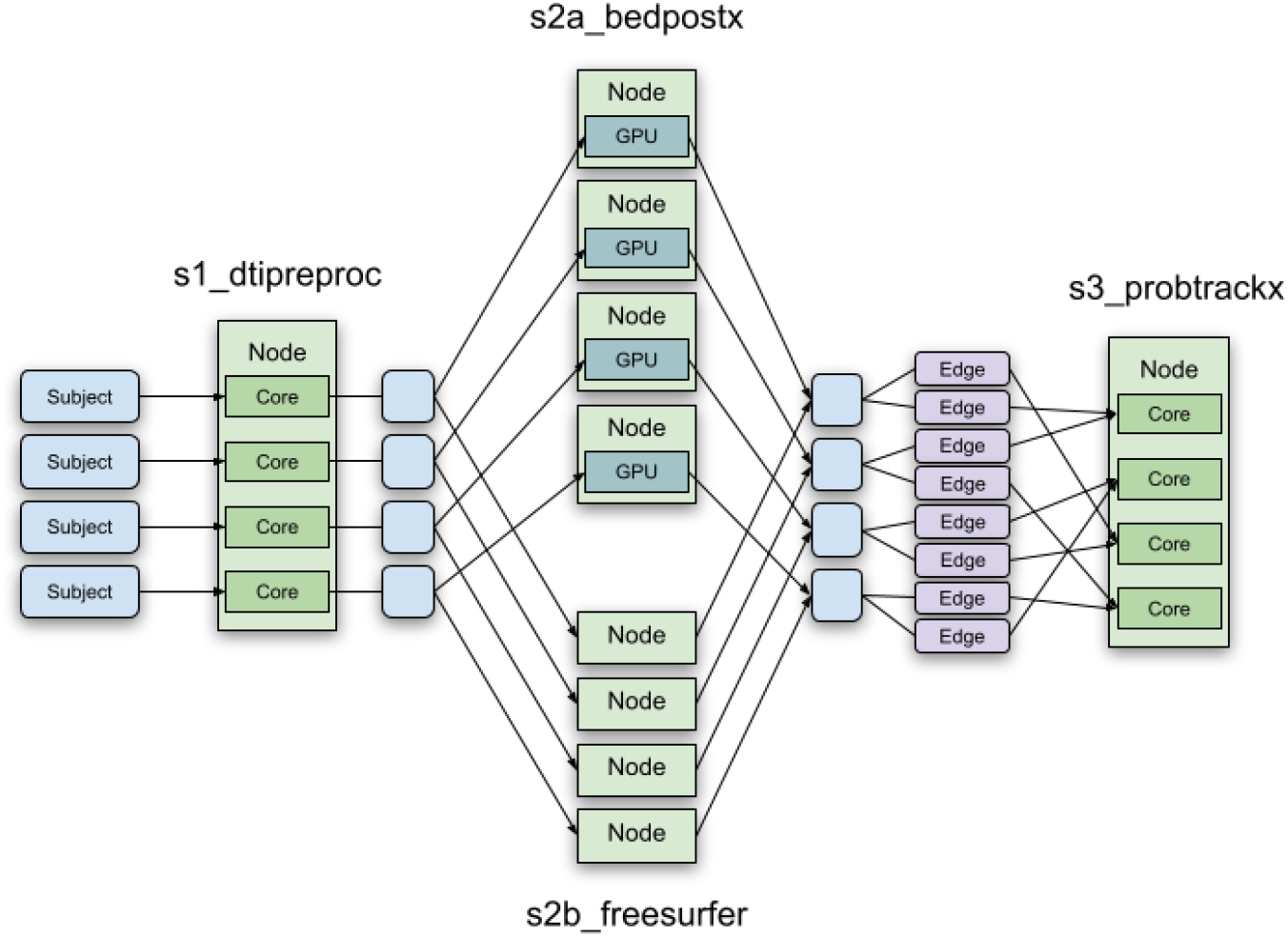
Example of Parsl allocation with heterogenous resources

**Figure 6:**
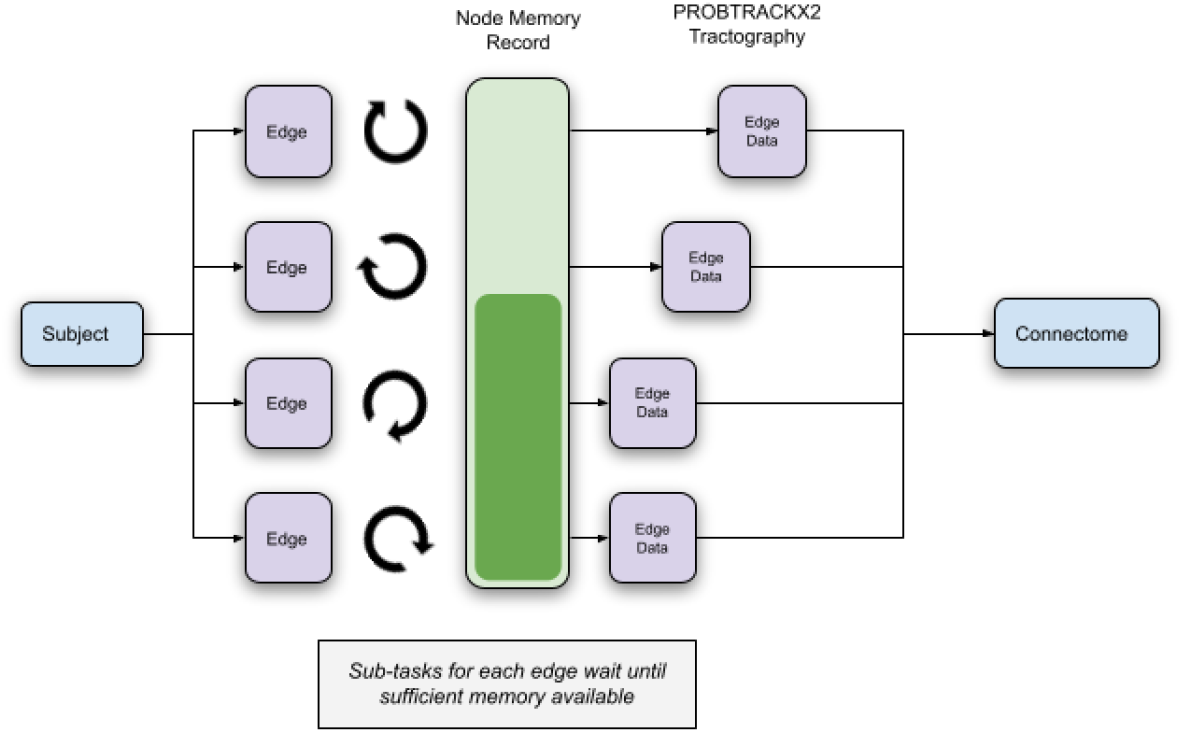
Memory management for tractography on a single subject

The original scripts come with several limiting characteristics. Exact versions of software libraries - FSL (Smith et al., 2004; Jenkinson et al., 2012), FreeSurfer (Fischl, 2012), and CUDA (NVIDIA et al., 2020) - must be installed locally. Input and output data must be stored at hard-coded paths. The user must specify resource allocation and submit batch jobs to the HPC scheduler manually. And parameters can only be adjusted by modifying the scripts themselves. Although the scripts are carefully tailored to run on specific HPC clusters, they are not sufficiently robust for anything more than experimentation with a handful of patients.

### 2.2. Subjects

Our new pipeline has been tested extensively on a large cohort of real-world patients. All participants were enrolled at eighteen Level 1 Trauma Centers across the USA as part of the prospective Transforming Research and Clinical Knowledge in Traumatic Brain Injury project (TRACK-TBI) (Palacios et al., 2020). TRACK-TBI is a National Institutes of Health–funded multi-center study that began in October 2013. The objective is to create a large, high-quality database that integrates standardized clinical, imaging, proteomic, genomic, and outcome measures to establish more precise methods for TBI diagnosis and prognosis.

TRACK-TBI patients were recruited after injury upon meeting the American Congress of Rehabilitation Medicine (ACRM) criteria for TBI. Other inclusion criteria were having a computed tomography brain scan as part of clinical care within 24 hours of injury, no significant polytrauma that would interfere with their assessment, and no MRI contraindications. Exclusion included prior major psychiatric or neurological pathology. Orthopedic trauma control patients were, like the TRACK-TBI patients, recruited from the Level 1 Trauma Centers, and presented mainly with lower extremity fractures. Orthopedic controls were ruled out for suspected head trauma, loss of consciousness, amnesia, previous TBI, or major psychiatric pathology. All eligible patients who voluntarily agreed to participate gave written informed consent. All study protocols were approved by the Institutional Review Boards of the enrollment centers.

MaPPeRTrac has been successfully run on 882 TRACK-TBI patients (median 30 yr; SD±15.0 yr; 184 female). This includes 152 patients from the pilot study (Yuh et al., 2014), 621 patients from the multi-site study (Palacios et al., 2020), and 109 patients from the orthopedic trauma control group (Bodien et al., 2018). Besides the orthopedic controls, all patients have been diagnosed with TBI. Patients were scanned at 1-2 weeks (mean 13.30 days; SD±2.10 days) after injury. The multi-site study and orthopedic controls were additionally scanned at 6 months (mean 184 days; SD±8.86 days) after injury. All MR imaging has been conducted with 3T scanners to generate whole-brain diffusion tensor images (DTI) with a variety of acquisition parameters. Further details can be found in the respective reference for each patient population (Yuh et al., 2014; Palacios et al., 2020; Bodien et al., 2018).

We henceforth refer to individual scans of patients as subjects. This results in a total of 1612 subjects processed with MaPPeRTrac. For performance testing, we have processed the 152 subjects from the TRACK-TBI pilot using the original scripts in addition to MaPPeRTrac. We stop at 152 subjects for economic reasons, since running the remaining 1460 subjects using the original scripts would be prohibitively expensive.

### 2.3. Compute Hardware

For testing performance of the pipeline, we used a compute cluster of nodes with Intel Xeon E5-2695 processors. Each compute node has 36 cores per node and a clock speed of 2.10 GHz (3.30 GHz with boost). All machines run the Tri-Lab Operating System Stack (LLNL, 2021) operating system and use Slurm scheduling. They have the following hardware characteristics.

**Table 1:** Hardware resources for Testing

| System | Pipeline Steps | RAM/Node | GPU |
| --- | --- | --- | --- |
| Quartz <sup>2</sup> | s1_preprocessing, s2b_freesurfer, s3_probtracx, s4_render | 128 GB | n/a |
| Pascal <sup>3</sup> | s2a_bedpostx | 256 GB | NVIDIA Tesla P100 |

Additionally, the pipeline has been successfully run on the following platforms to demonstrate portability.

**Table 2:** Additional Tested Platforms

| System | CPU | OS | Scheduler | RAM/Node | GPU |
| --- | --- | --- | --- | --- | --- |
| Catalyst <sup>4</sup> | Intel Xeon E5-2695 | TOSS 3 | Slurm | 128 GB | n/a |
| Wynton <sup>5</sup> | Intel Xeon E5-2640 | CentOS 7 | Grid Engine | 48-512 GB | NVIDIA GTX 980 Ti |
| Blues <sup>6</sup> | Intel Xeon E5-2670 | CentOS 7 | Slurm | 768 GB | NVIDIA Tesla K40m |
| Cooley <sup>7</sup> | Intel Xeon E5-2620 | Cray Linux | Cobalt | 384 GB | NVIDIA Tesla K80 |
| Amazon <sup>8</sup> | Intel Xeon (Skylake vCPU) | Ubuntu | AWS | 61 GB | NVIDIA V100 |

## 3. MaPPeRTrac

MaPPeRTrac accomplishes the goals of the original tractography scripts, but with a significantly faster, parallel, more portable, and better parameterized implementation. It takes advantage of parallelism at both the subject-level and task-level coupled with GPU acceleration in order to significantly speed up connectome generation on HPC clusters. It also pre-processes raw DICOM files from a wide variety of MRI scanners, dynamically handles resource allocation and parameter configuration, and performs heuristic validation on intermediate data outputs. This level of complexity and robust execution is managed by a clear functional hierarchy and containerization of software dependencies, ensuring that users can easily modify and deploy the tools on any HPC cluster.

The specific software commands used by MaPPeRTrac are intended to be identical to those used by the original scripts. Further details can be found in (Owen et al., 2015, 2016) or Appendix B. However, we are more interested in the architectural features that enable these commands to run at scale and ease usability.

### 3.1. Pre-processing

One of the limitations of the original scripts is their inability to read data directly from MRI scanners. Converting raw DICOM files into a diffusion tensor image (DTI) and T1-weighted anatomical volume is a laborious process, even with automation tools. This is made more challenging by the wide variety of MRI scanners and file standards from major manufacturers, specifically GE, Philips and Siemens. MaPPeRTrac overcomes this obstacle by implementing a novel algorithm to universally pre-process DICOM files. It converts all DICOM files into NIfTI format and applies a series of statistical heuristics to determine whether to process them into the DTI, anatomical volume, or b-value weighting. This capability, coupled with adoption of BIDS standards, enables our pipeline to handle inputs regardless of naming convention or file structure.

At times, pre-processing MRI inputs without human oversight runs the risk of faulty behavior. Additional heuristics in the pipeline help catch invalid data. The pipeline alerts users if data are clearly missing or significant outliers are found. After manually reviewing a random sample of 200 subjects processed directly from DICOM files, we have found none that are handled incorrectly by this pre-processing step. Nor do any graph analyses of the 1612 connectomes we have generated show any indication of faulty pre-processing.

### 3.2. Parallelization

The original scripts for generating structural connectomes are a monolithic sequence of commands. We modularized the components of the pipeline and implemented parallel execution of the modules. In our efforts to modularize and parallelize the different analysis steps, we leveraged a parallel scripting language called *Parsl* (Babuji et al., 2019) that is being developed at University of Chicago and is designed to process massive batches of data on heterogeneous computing resources. *Parsl* is especially designed to work seamlessly on various DOE computational resources. The modularization of various analysis steps helped profile various modules where we measured the computational (CPU, memory, I/O) requirements of individual steps, gain insights into parallelization possible by virtue of tracking data dependencies among different stages of the pipeline and helped define and tailor resource requirements for specific steps. Previous neuroimaging pipelines tend to reserve maximum computational resources available which remains unchanged throughout the execution of the pipeline, so that the most expensive steps can complete. This strategy results in sub-optimal utilization of computational resources and typically disallowed shared computational facilities. To address this challenge, we implemented a dynamic allocation of the computational resources required at the time of a particular task execution based on the computational profile of the task and relinquishing resources when they are no longer required.

*Parsl* defines the pipeline architecture using the data dependencies among the steps and dispatches tasks to the underlying schedulers tasks that have all the input datasets available. This feature enabled MaPPerTrac to exploit parallelism at pipeline level and at the individual task level. For example, if there are 1000 MRI input datasets available to process, all of the samples can be submitted at once and the pipeline will request resources to process the first step of the pipeline for all the samples in parallel. Subsequent steps of the pipeline are scheduled appropriately when the inputs of the steps are generated. In addition to pipeline-level parallelization, we implemented task level parallelization where tasks in the pipeline that do not have data dependencies get scheduled and executed in parallel. In MaPPeRTrac, this is manifested in parallel execution of parcellation and tensor estimation on the same subject.

The pipeline also takes advantage of sub-task level parallelization where we exploit the multiple CPU cores present in a compute node and perform the computation in parallel. This is achieved by dividing the input data into smaller chunks and running the analysis on individual CPU cores. Whereas sub-task level parallelization is implemented in FreeSurfer using OpenMP (Dagum and Menon, 1998), it is achieved with pre-processing and tractography using *Parsl*’s job allocation system.

### 3.3. Parameterization

The implementation of the pipeline using *Parsl* involves creation of a directed acyclic graph where the edges are data dependencies among the steps. Since all the steps in the pipeline are run without user intervention, the execution engine needs to have all the parameters for all the steps be available. The need for parameterization of individual steps is even greater when executing a large batch of the MRI datasets as it is impractical to have a human operator check the outputs of a step and launch the subsequent step manually. Since the graph of tasks is specified when launching *Parsl*, this architecture necessitates parameterization of the elements required to describe the tasks and sub-tasks. All the pipeline parameters are configurable from a single source, which can either be commandline arguments or a configuration file. When left unspecified, the pipeline will estimate appropriate parameters or use default values from the original script collection. This enables users to easily orchestrate complex jobs across different HPC clusters.

### 3.4. Portability

The *Parsl* framework is implemented in Python, one of the most popular cross-platform programming languages. The pipeline is implemented as a Python package so it can be used on any platform that supports Python. One of the strengths of the *Parsl* framework is the separation between definition of tasks and the actual execution of the tasks. Once the tasks are defined in *Parsl*, its plug-in based architecture enables execution on most of the modern HPC schedulers.

*Parsl* supports execution on computational clouds, supercomputers and HPC clusters seamlessly using plugins for various computational elements. Each provider implementation may allow users to specify additional parameters for further configuration. Parameters are generally mapped to resource manager submission script or cloud API options. Examples of local resource-manager-specific options are partition, wall clock time, scheduler options. This can include scheduling headers such as #SBATCH for Slurm (Yoo et al., 2003) or worker initialization commands (e.g., loading a Conda environment). Cloud parameters include access keys, instance type, and spot bid price. At the time of writing this manuscript, *Parsl* supports 12 *providers* that include the gamut of frameworks that deliver computational resources. To overcome the differences in these compute elements, and present a single uniform interface, *Parsl* implements a simple provider abstraction. This abstraction is key to its ability to enable scripts to be moved between resources. The provider interface exposes three core actions: submit a job for execution (e.g., sbatch for the Slurm resource manager), retrieve the status of an allocation (e.g., squeue), and cancel a running job (e.g., scancel). *Parsl* implements providers for local execution (fork), for various cloud platforms using cloud-specific APIs, and for clusters and supercomputers that use a Local Resource Manager (LRM) to manage access to resources, such as Slurm (Yoo et al., 2003), HTCondor (Thain et al., 2005), and Cobalt (Desai, 2005). By leveraging *Parsl* to define our pipeline, we have been able to achieve portability of execution.

After achieving execution portability using *Parsl*, we have examined different ways in which we can create portable bundles of applications that can be packaged and used in conjunction of the pipeline. Examining past work in reproducible neuroscience such as (Theaud et al., 2020), we have concluded that containerization technologies are key to ensure portability. Software containers are portable execution elements that provide an abstraction over underlying operating system and hardware to to improve the portability of applications. Containerization technologies are supported by computational clouds, DOE supercomputers, and campus HPC providers. Two popular containerization technologies exist with important technical differences. In order to support extensive portability, we generated containers using both *Singularity* (Kurtzer et al., 2017) and *Docker* (Boettiger, 2015) and made these containers publicly available. In addition to increasing the portability of the pipelines, the containerization technologies also enable versioning of various analysis tools that make up the pipeline. New versions of the analysis tools, for example, a new version of FreeSurfer, can easily be tested by updating the ‘recipe’ for generation of the container to point to the location of the updated version of FreeSurfer. Our recipes for *Docker* and *Singularity* containers contain all the tools and their dependencies for successful execution of the analysis. Users can either download an existing pre-built container or use the recipes to build it themselves. This is particularly important since a major hurdle to scientific software development is the onerous installation of tools and their dependencies.

The addition of numerous features and performance improvements to the pipeline reflects commensurate growth in the complexity of its software and output data. To mitigate complexity, we strictly adhere to the Brain Imaging Data Structure (BIDS) (Gorgolewski et al., 2016), a protocol for standardizing neuroimaging data. The DICOM and NIfTI inputs are organized and shared along with tractography outputs, ensuring datasets across the entire pipeline remain transparent. Adoption of the BIDS data format addresses another key portability challenge of how the inputs are described and where the outputs of various steps of the pipeline are available.

### 3.5. Performance enhancements and other improvements

In addition to the performance improvements resulting from node-level and core-level parallelization of the pipeline, we have made significant improvements in specific tools.

#### 3.5.1. Dynamic memory management of PROBTRACKX2

We use PROBTRACKX2 for performing probabilistic tractography, which is very memory-intensive. Though *Parsl* has a wide range of features to manage resource allocation, the configuration and optimization of individual tasks has to be done using the computational profiling of the tool using different inputs. This has been challenging as the computational (especially memory) requirements vary across different datasets.

Because we generate the inputs to PROBTRACKX2 at run-time, it is difficult to predict just how much memory each task will need. Overestimating memory usage prevents parallelization entirely, because the worst-case tasks can consume an entire node’s memory. But if we underestimate memory usage, parallel tasks will request out-of-bounds addresses and invariably crash (even with strict paging on the computing hardware). This appears to be the result of PROBTRACKX2’s original design, which is not intended to run in parallel on the same node. In order to run it safely in parallel, we estimate each task’s memory usage at run-time and record the total estimate for each node in a thread-safe file. New tasks attempting to launch PROBTRACKX2 will wait until their node’s estimated memory usage falls below a safe threshold, found in testing to be between 50 to 70 percent of the node’s total memory. After completing a PROBTRACKX2 run, each task updates the record to free up memory and allow other tasks to proceed.

#### 3.5.2. Performance tracking code

We enhanced the pipeline by computational profiling code that measures core-time and wall-time of each step for each subject. The pipeline saves performance data and logs to the file system after each task is completed. Since each parallel process records data independently, we were able to collate performance data into a global timing log. This log, which contains resource usage and wall-clock time for each processing step, enables users to make informed estimates in future computations.

#### 3.5.3. Automatic visualization

The final step of the pipeline is to render EDI generated by the pipeline. Image rendering is even possible on headless nodes as we included the VTK library (Schroeder et al., 2004) with the container thus increasing the usability of the pipeline out of the box. However, this step is disabled by default. This is because it adds complexity that may discourage users, especially as most researchers analyze neuroimaging data using interactive tools such as FSLeyes instead (Jenkinson et al., 2012). For additional details see Appendix B.

Enhancements that did not sufficiently improve performance or usability are outlined in Appendix D.

## 4. Results and Evaluation

**Table 3:**
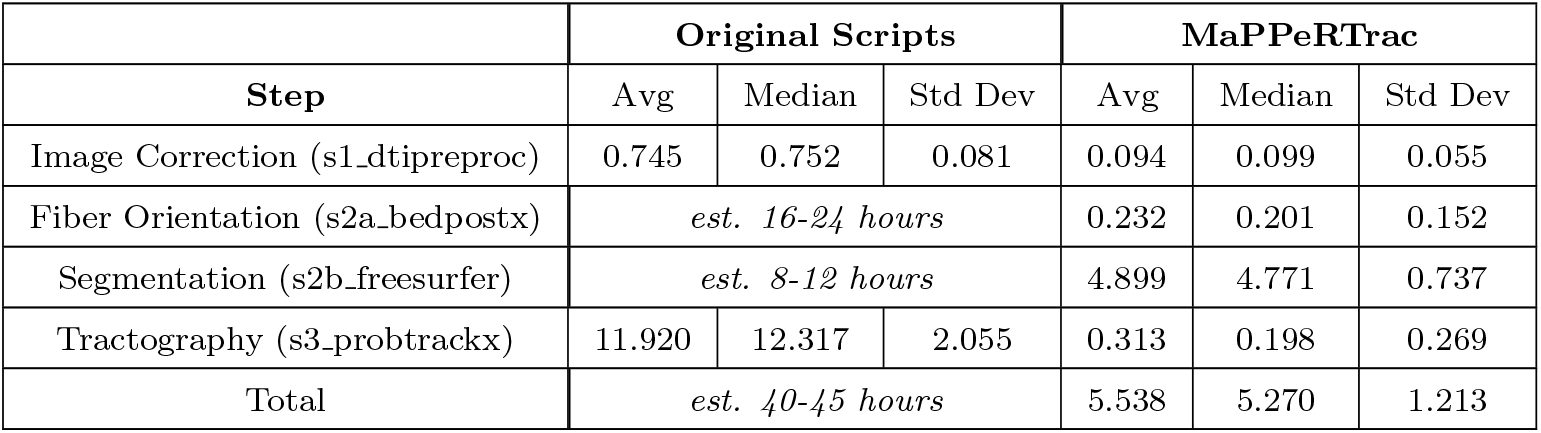
Detailed Wall Time (hours)

**Table 4:** Detailed Core-Hours

| Step | Original Scripts |  |  | MaPPeRTrac |  |  |
| --- | --- | --- | --- | --- | --- | --- |
|  | Avg | Median | Std Dev | Avg | Median | Std Dev |
| Image Correction (s1_dtipreproc) | 13.410 | 7.886 | 3.406 | 0.895 | 0.360 | 0.685 |
| Fiber Orientation (s2a_bedpostx) | *est. 720 core-hours* |  |  | 9.283 | 7.793 | 3.443 |
| Segmentation (s2b_freesurfer) | *est. 360 core-hours* |  |  | 191.294 | 199.336 | 8.306 |
| Tractography (s3_probtrackx) | 429.120 | 339.853 | 158.012 | 30.962 | 31.734 | 1.322 |
| Total | *est. 1500 core-hours* |  |  | 5.538 | 5.270 | 1.213 |

### 4.1. Performance

Compared to the original tractography scripts, we have achieved enormous speedups at all stages of tractography. Figure 7 shows improvements to the average time to compute a single connectome by MaPPeRTrac versus the original scripts, broken down by each step. In the most dramatic case, fiber orientation runs 8600% faster, thanks to GPU acceleration. Image correction and tractography also see significant speedup by spreading sub-tasks across nodes. Segmentation has a more modest speedup, since it can only distribute processes across cores on a single node. The average total speedup for a single subject is 735%, from 40 to 5.5 hours.

**Figure 7:**
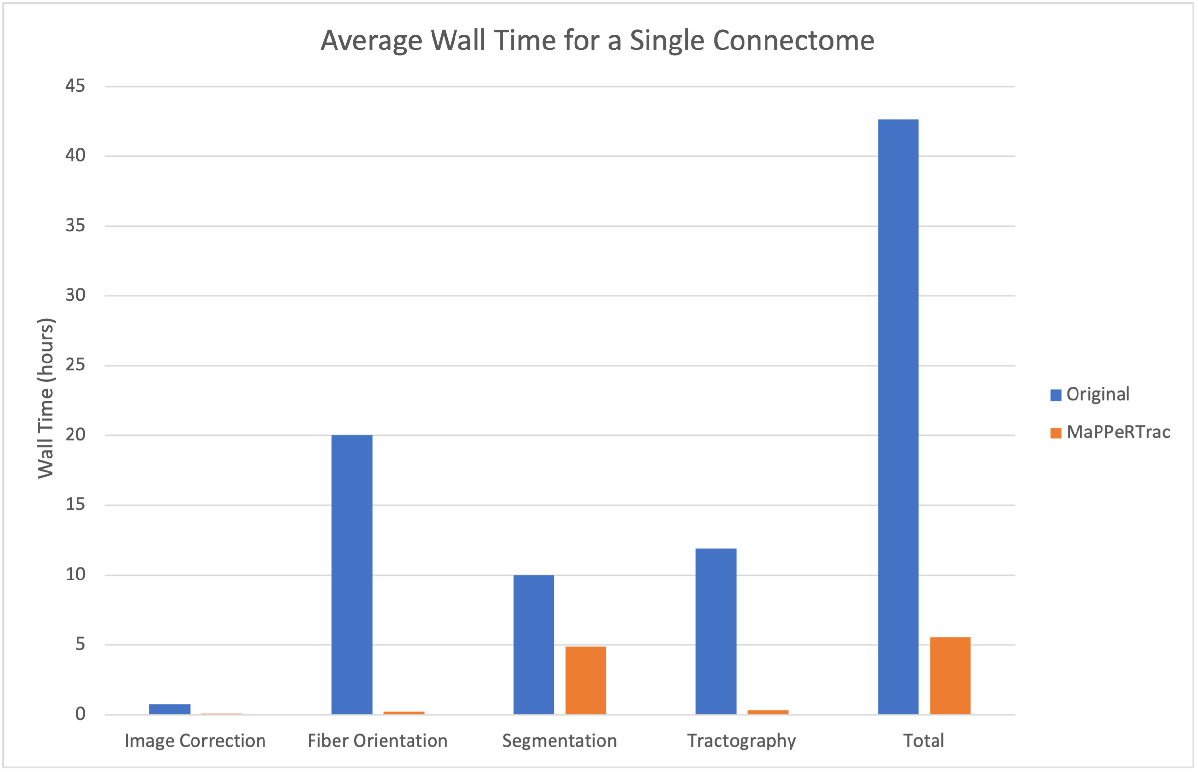
Average time for a single connectome

**Figure 8:**
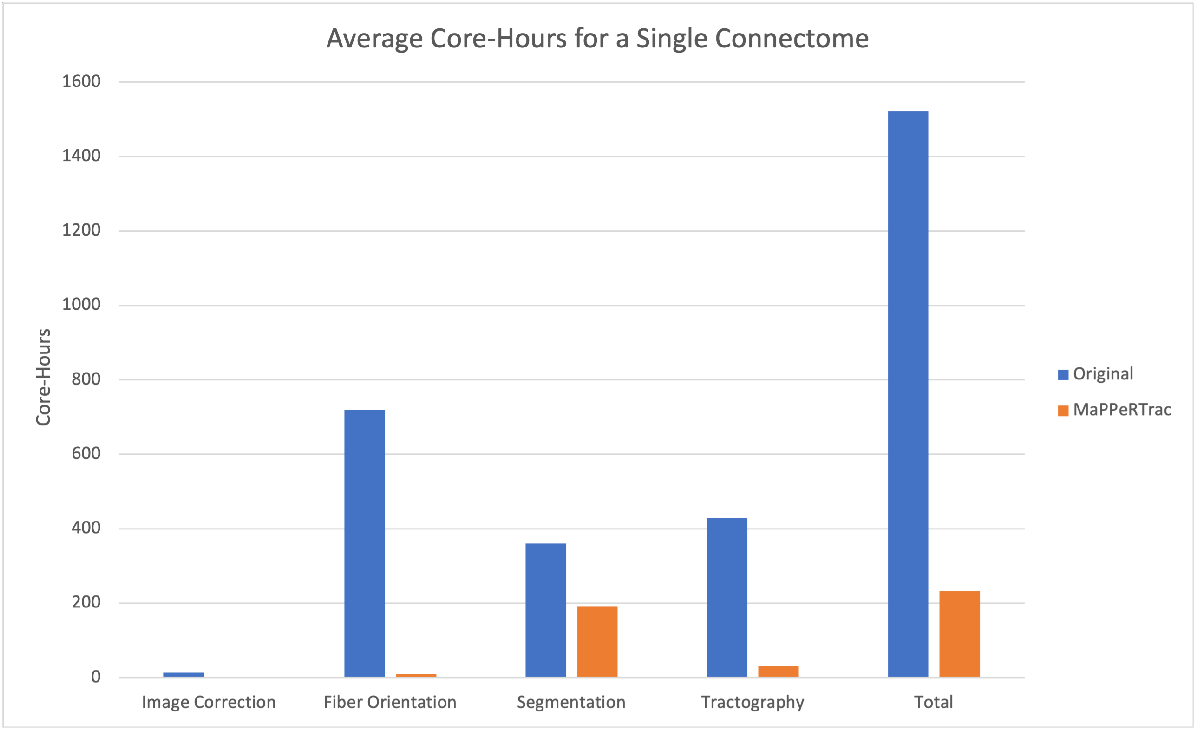
Average resource consumption for a single connectome

For resource-constrained users, a more practical metric is average core-hours. This measures the average compute time for a single connectome multiplied by the number of utilized computational cores. Since much of our performance gains are derived from parallelization, it is necessary to compare how much of this improvement derives from resource availability alone. In particular, tractography consumes a vast amount of core-hours that can only be mitigated by parallelization across many nodes. Nevertheless, MaPPeRTrac on average consumes a total of 625% fewer core-hours.

As explained in 2.2, we have used a subset of 152 subjects from TBI patients for performance testing. However, even on this subset, processing the original scripts’ implementation of fiber orientation and segmentation would be prohibitively expensive. Therefore, those particular values (e.g., *est. 16-24 hours*) reflect in-depth interviews with neuroscientists who use the original scripts extensively. Since our primary goal is to produce an efficient tractography pipeline, we have chosen not to waste vast resources on redundant computation and instead rely on expert testimony.

### 4.2. Parameterization

As described in 4.2, we have implemented a powerful parameterization framework that can use either command-line arguments or a JSON configuration file. We include several examples of configuration files for parameterized execution on various computational platforms in Appendix A.

### 4.3. Portability

The pipeline has been running at two supercomputing resources and a university computing cluster with minimal site-specific changes that were made to a single configuration file. The site-specific changes include the location of input files and the bespoke user options that *Parsl* needs to interact with job schedulers. When executing on a cloud, the configuration options also include account information and location of security credentials. These configuration elements do require a certain technical know-how, which may be a barrier to entry.

Another limitation is maintaining the latest software packages in the *Singularity* container. Although MaPPeRTrac has been designed to be as self-encapsulated as possible, hardware and driver support may fall out of date. Support for different versions of NVIDIA CUDA is especially challenging, since newer hardware tends to break compatibility with older drivers. Updates to *Parsl* may also break compatibility since MaPPeRTrac’s Python scripts are not containerized.

Nevertheless, our collaborators at multiple institutions have validated the portability and ease-of-use of our pipeline by running it with growing independence. For instance, the pipeline has run successfully on the Wynton HPC platform at the University of California, San Francisco for a test set of TRACK-TBI data, and produced expected outputs from all steps of the pipeline that were consistent with outputs from LLNL and ANL platforms. The *Parsl* framework and our pipeline were robust to accommodate computing environments, such as Wynton, that use the Sun Grid Engine scheduler (Gentzsch, 2001), which is less commonly supported compare to Slurm or HT-Condor (Tannenbaum et al., 2001). Additional configuration details for Wynton and Sun Grid Engine are detailed in Appendix C.

### 4.4. Enabling FAIRness

In line with our objectives, the pipeline embodies and enables the FAIR principles. To ensure the pipeline’s artifacts remain Findable and Accessible, we published the source code to GitHub with a permissive BSD license. We also uploaded the container recipe to Singularity Hub, the official registry for the *Singularity* software library. Furthermore, the pipeline accepts input identifiers from sources such as OpenNeuro. It generates similar identifiers along with the output artifacts. None of the input, output, or intermediate files are proprietary – the pipeline uses BIDS-compliant open source formats such as NIfTI. We standardized inputs and outputs of the pipeline according to the BIDS format. DICOM sources, NIfTI images, and pipeline derivatives are kept in separate folders, organized according to their patient and session. This makes it easy to compare different timepoints and retests, since they are always located together.As a result, the pipeline can provide a clear provenance to all of its digital outputs.

### 4.5. Conclusions and Future Work

Progress in connectomics has been limited by steep computational cost of probabilistic white matter fiber tractography, complexity in installing applications and dependencies, and challenges in scaling while adhering to best practices for reproducible research. We have developed MaPPeRTrac to enable high performance, parallel, parameterized, and portable generation of connectomes that is well-tested, robust and easy to use for the community. To lower the barrier to entry for users, we ultimately plan to make the pipeline available as a service on public cloud computing resources so that researchers can upload data and generate connectomes using the state-of-the-art tools without becoming a computational expert.

## Appendix A. Examples of MaPPeRTrac Parametrization

1. Quartz (LLNL)
2. Cooley (ANL)
3. Blues (ANL)
4. Amazon Web Services

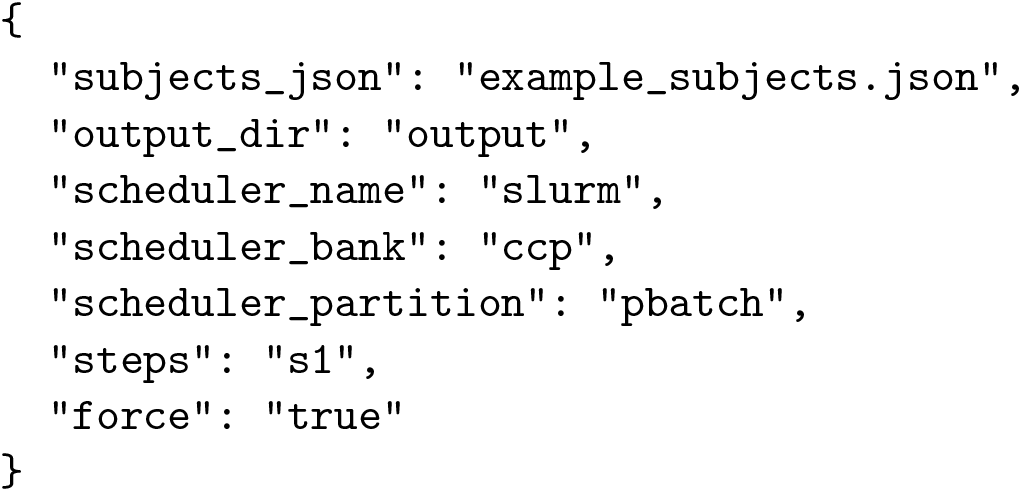

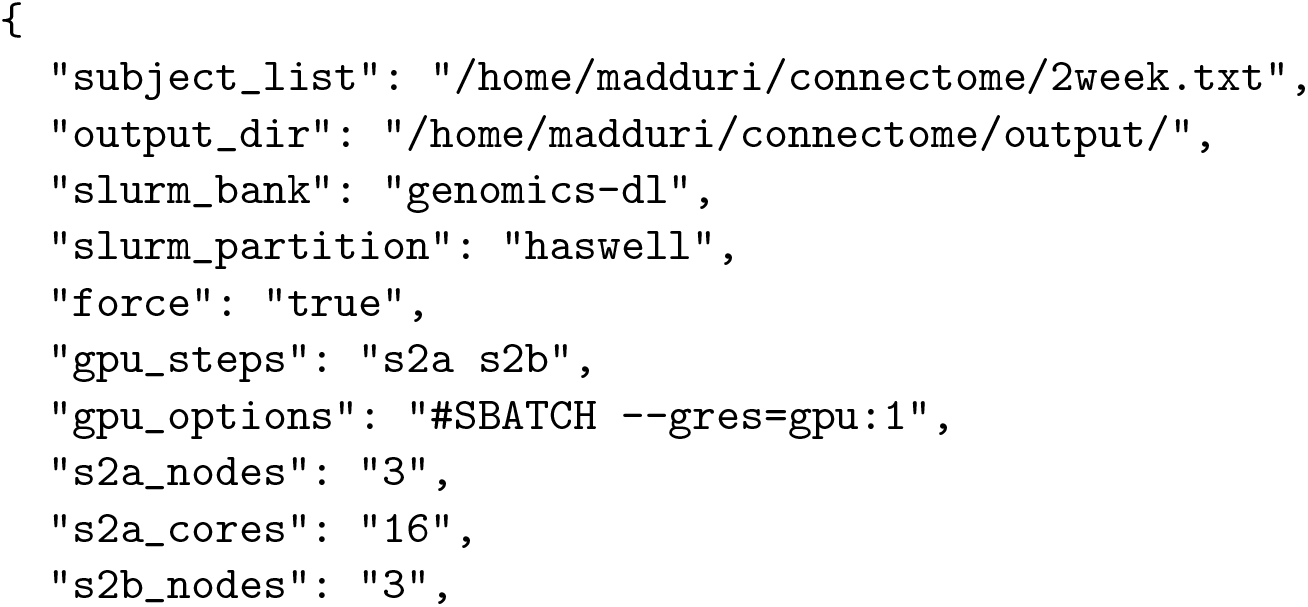

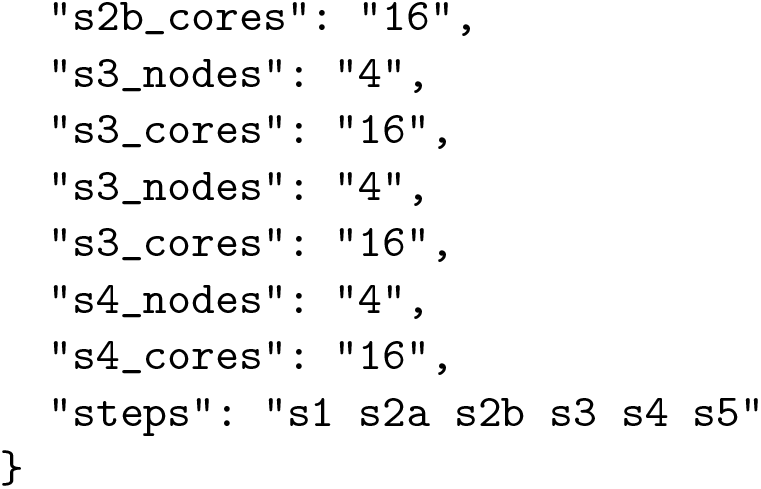

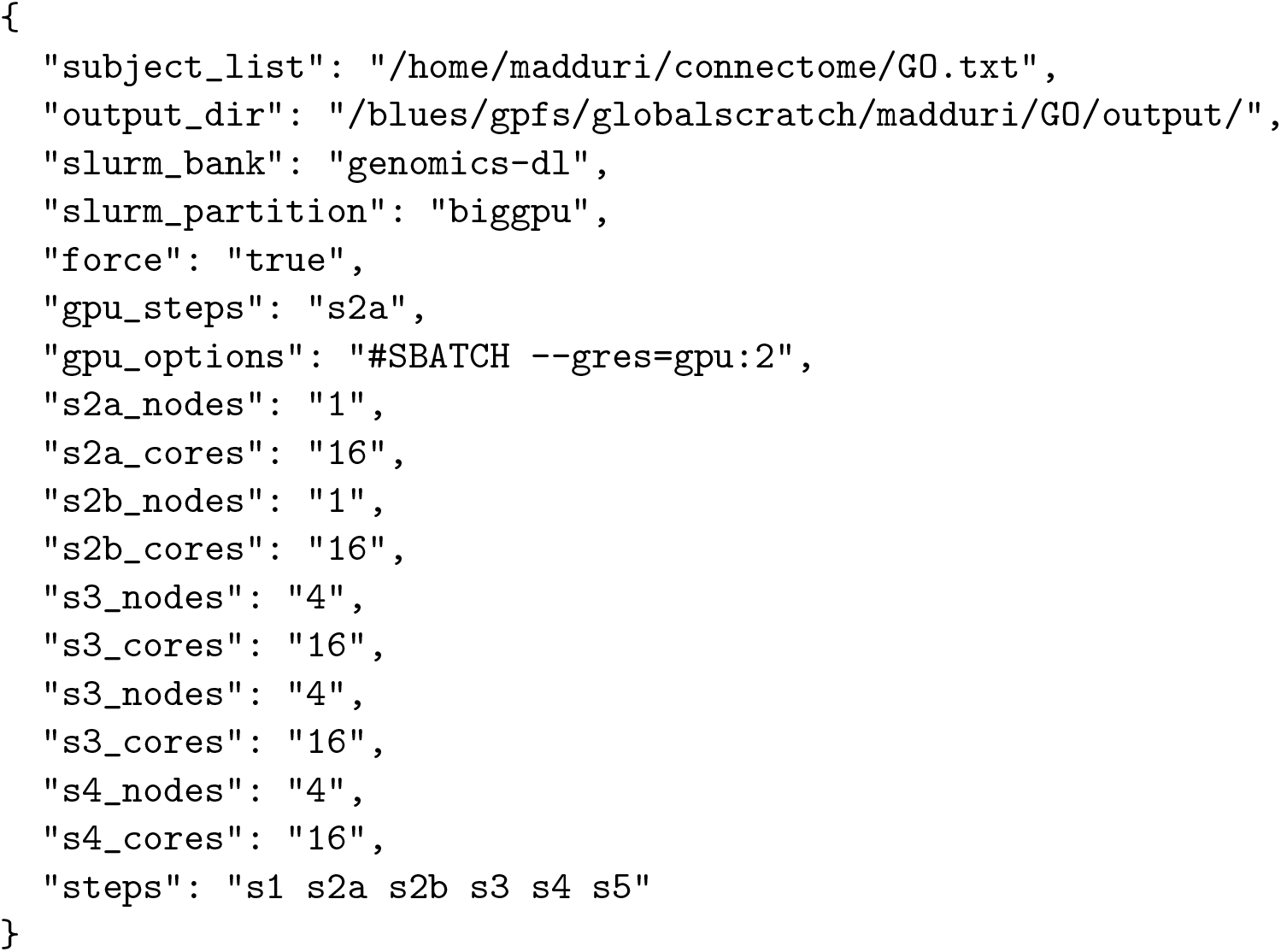

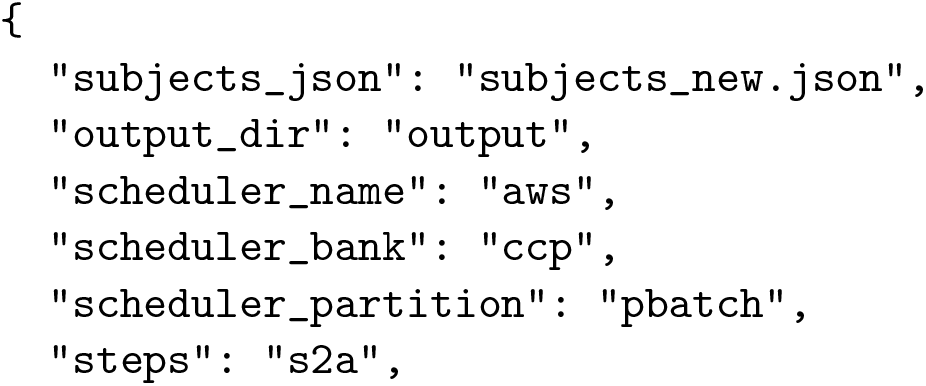

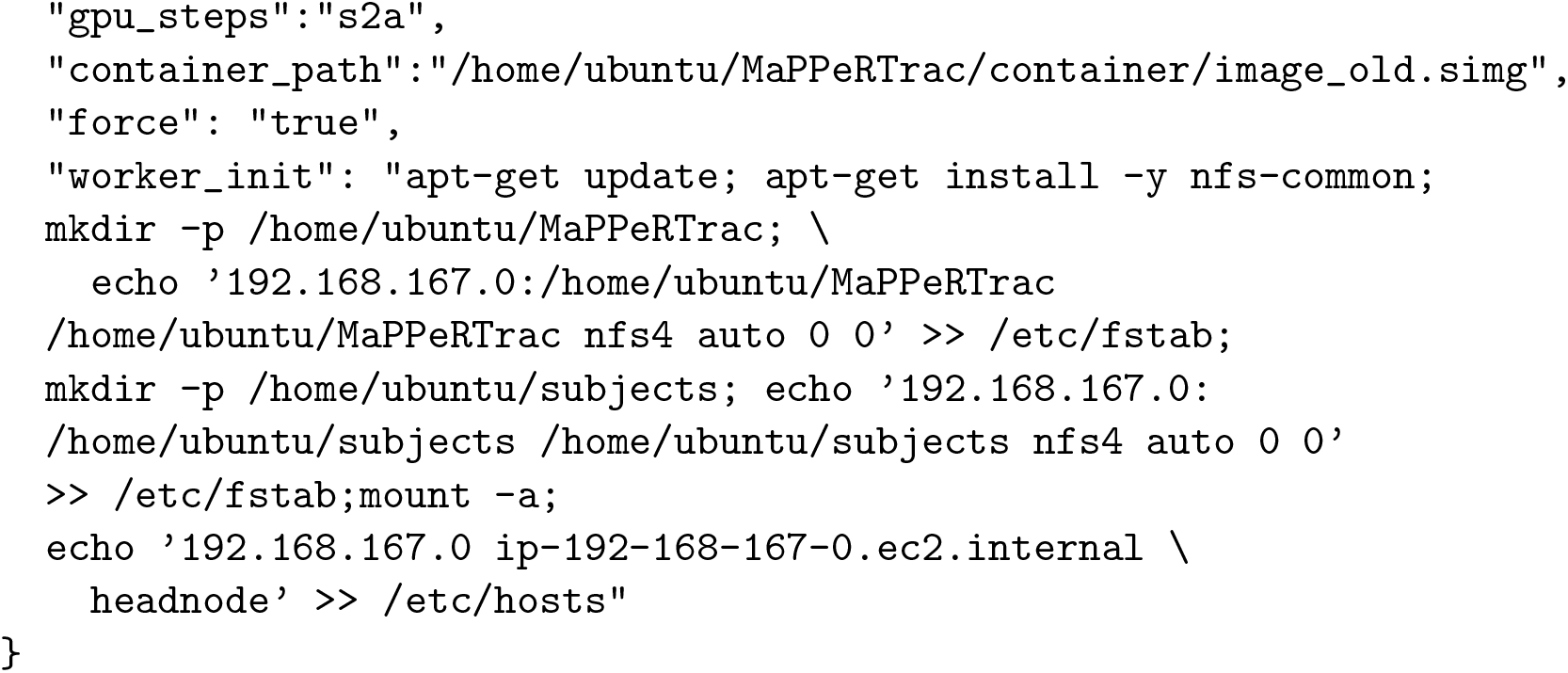

## Appendix B. Neuroimaging Software Details

The pipeline and instructions to the run are available from GitHub (https://github.com/LLNL/MaPPeRTrac). The pipeline divided into steps, which can be run individually or sequentially

1. *s1 dti preproc.py* – this step corrects motion, eddy current, and noise. The Brain Extraction Tool (BET) is used to isolate the brain from surrounding tissue. Images are corrected for motion and eddy currents using the FMRIB linear-image registration tool (FLIRT) with a 12-parameter linear image registration using the b = 0 s/mm2 image as the reference. Using FLIRT, the FA map of every patient and session is registered to the T1 in order to obtain a diffusion to structural transform.
2. *s2a bedpostx.py* - estimates the fiber orientation at every voxel using BEDPOSTX with default settings.
3. *s2b freesurfer.py* - performs cortical parcellation using FreeSurfer with the Desikan–Killiany atlas, resulting in 68 cortical regions and 14 subcortical regions. The 68 cortical regions are transformed to the gray–white matter boundary (GWB). These 82 regions represent the nodes of the connectome. Additionally, we register the FA volume to the T1 volume. Each of the cortical GWB volumes and the subcortical volumes are registered to the diffusion space to be used as seeds for the tractography. Steps s2a and s2b can be run in parallel.
4. *s3 probtrackx.py* – runs probabilistic diffusion tractography, generating connectomes and EDI. We use PROBTRACKX2 to generate 1000 streamlines from each seed voxel. The target region is used as a way-point mask and all other regions excluded. Given 82 regions in the D-K atlas, this results in 82*82 - 82 = 6642 tractography runs per subject. Tractography results are binarized to create a mask of white matter voxels needed to connect each pair of cortical/subcortical regions. This uses a consensus connectome based on (Raji *et al.*, 2020) to generate the final connectome and EDI.
5. *s4 render.py* – runs the VTK render suite on EDI outputs using a copy of vtkpython bundled in the Singularity container. We render a scaled average of edge density as well as slices along the horizontal and sagittal planes. In order to make all renders visible, we normalize the edge density between 0 and 1. The minimum and maximum edges per voxel are written at the bottom of the image, so that users can understand the actual density presented.

**Figure B.9:**
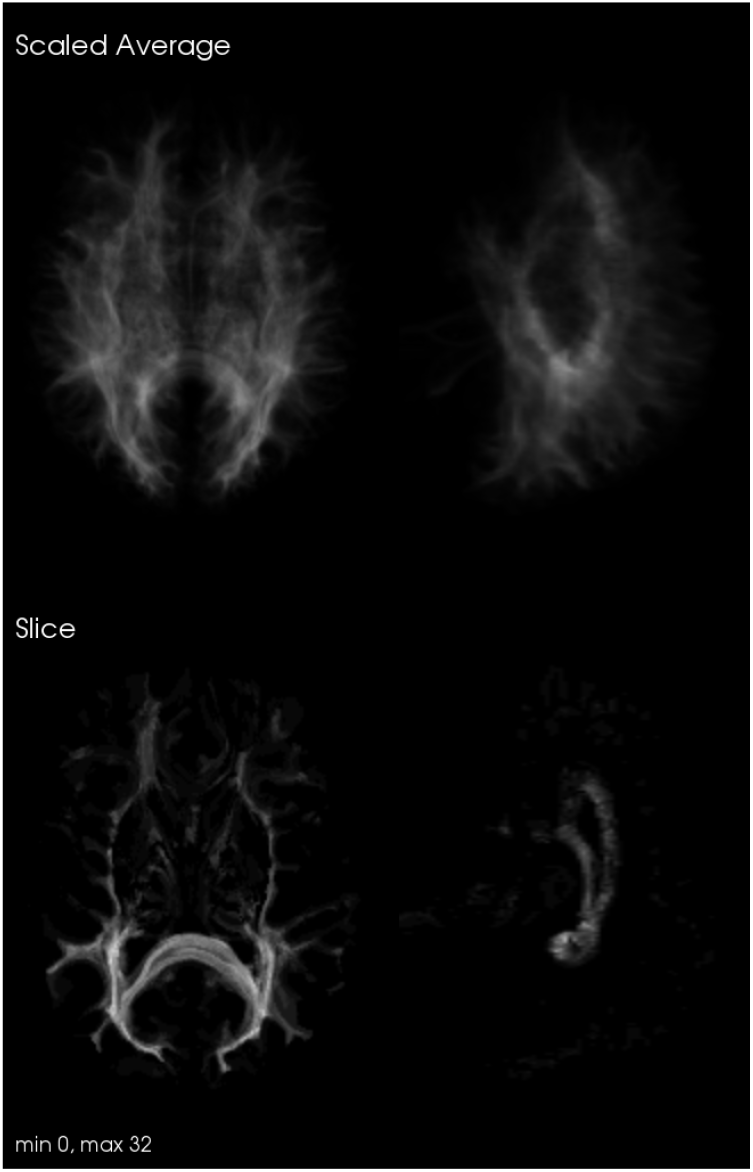
Example of visualization generated by s4 render

## Appendix C. Configuration of MaPPeRTrac on the “Wynton” cluster using Sun Grid Engine

In our efforts to test portability of the pipeline across multiple HPC schedulers, we discovered some gaps in the support for the soon-to-be-deprecated Sun Grid Engine. We worked with the Parsl team to close these gaps and improved the overall portability of the pipeline and aided development of Parsl’s support for SGE scheduler. We configured SGE as a Parsl *provider* in the following way:

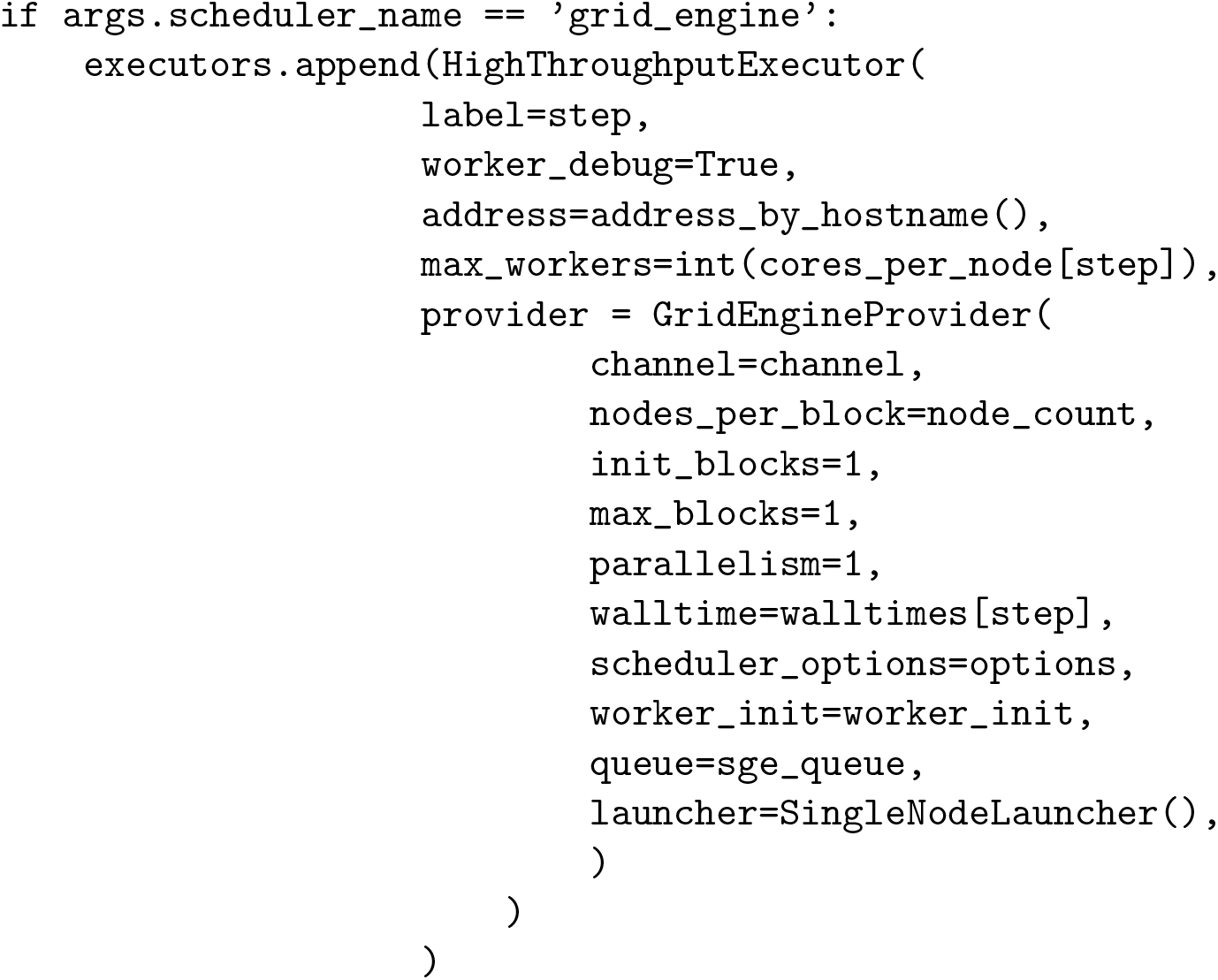

Specifically, “*queue*” was added to the latest version of Parsl upon our request. This parameter enabled us to specify the name of the queue on Wynton to which MaPPeRTrac submits the *parsl.sge* jobs resulting from execution of the various pipeline steps. The job queue specification was necessary for steps intended to run on GPU nodes (scheduled via *gpu.q*), otherwise *parsl.sge* jobs would be defaultly assigned to CPU nodes (scheduled via *member.q*, *short.q*, or *long.q*). The *queue* specification parameter for other schedulers has been supported by Parsl since its earlier versions but wasn’t available for SGE until Parsl approved our recent modifications to the Parsl source code repository.

Another parameter specific to SGE was the *max workers* for the *High-ThroughputExecutor* configuration. *HighThroughputExecutor* sets *max workers* to infinity by default. We configured this parameter to be consistent with the allowance of computing resources on Wynton. The decision on the exact value of *max workers* comes from a trade-off between job priority and requested computing resource. Requesting too many workers (a.k.a. threads) for one job can significantly impact the waiting time before the job gets running, but requesting too few cores limits the running speed. We have been using up to 72 *max workers* for our testing jobs.

The testing set of data at University of California, San Francisco consists of diffusion MRI scans in anonymized NIfTI format from three subjects in the patient group of the TRACK-TBI study. For a single subject, s1 preprocessing took 4 minutes on 1 CPU node with 72 threads; s2a bedpostx took 18 minutes on one GPU; s2b freesurfer took 4 hours 16 minutes on 1 CPU node with 16 threads; and s3 probtrackx took 1.5 hours for a collection of 930 edges that was split into 10 lists and submitted to 10 GPUs, respectively. The FSL probtrackx2 gpu computation for each edge took an average of 36 seconds on a GPU. Taking into account that s2a and s2b have no data dependency and are capable to run simultaneously, the total run time for one subject is approximately 6 hours. This run time performance is closely comparable to the performance on Quartz at LLNL (5.5 hours).

1. Configuration for s1 preprocessing.py
2. Configuration for s2a bedpostx.py
3. Configuration for s2b freesurfer.py
4. Configuration for s3 probtrackx.py

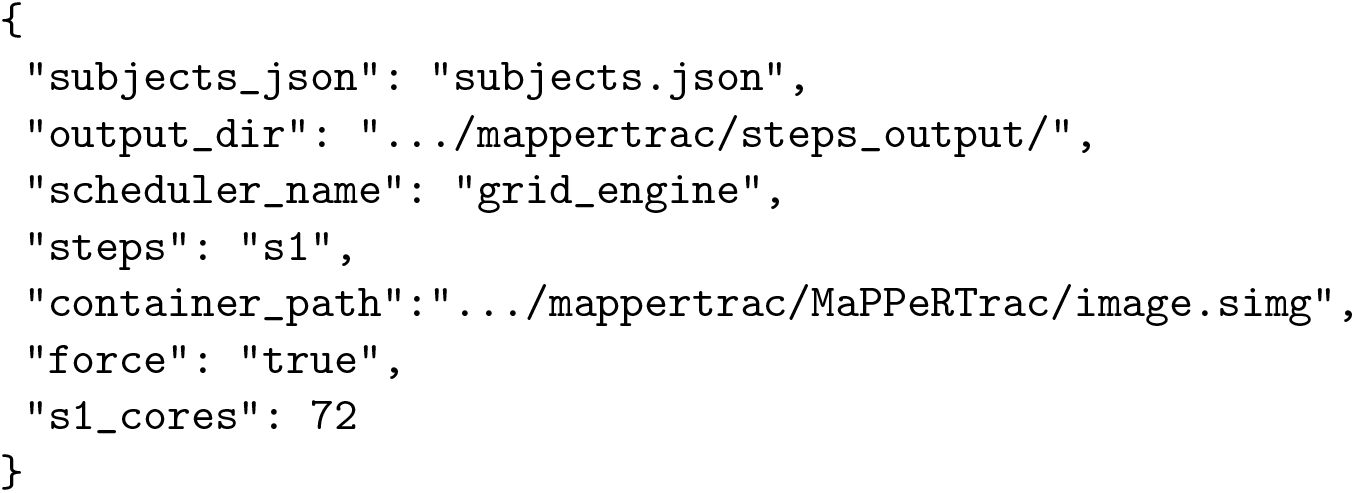

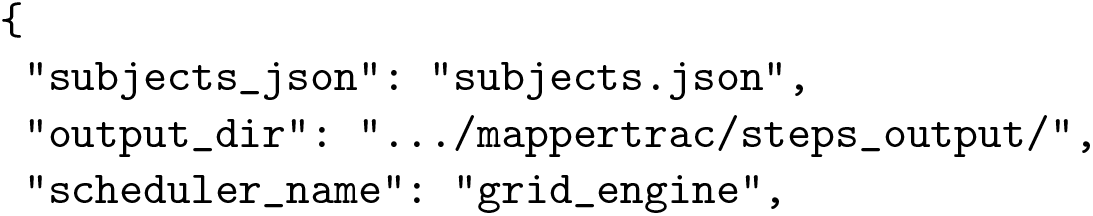

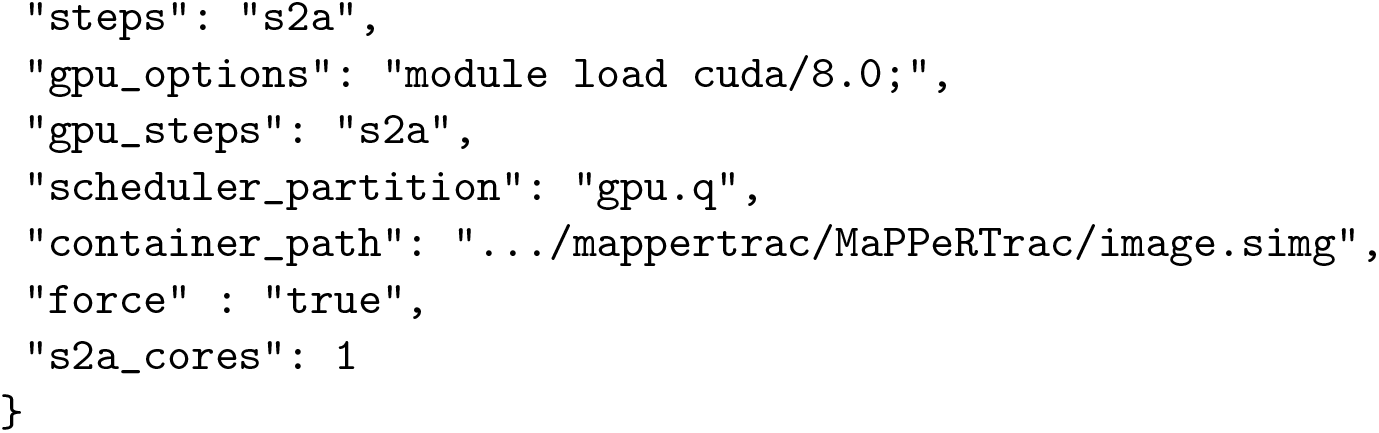

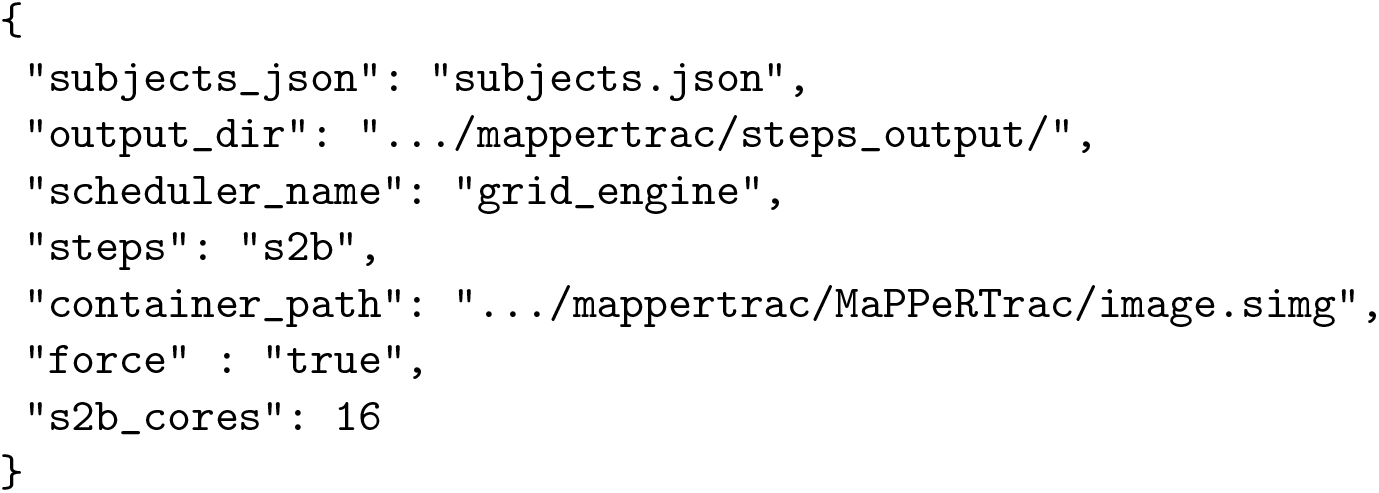

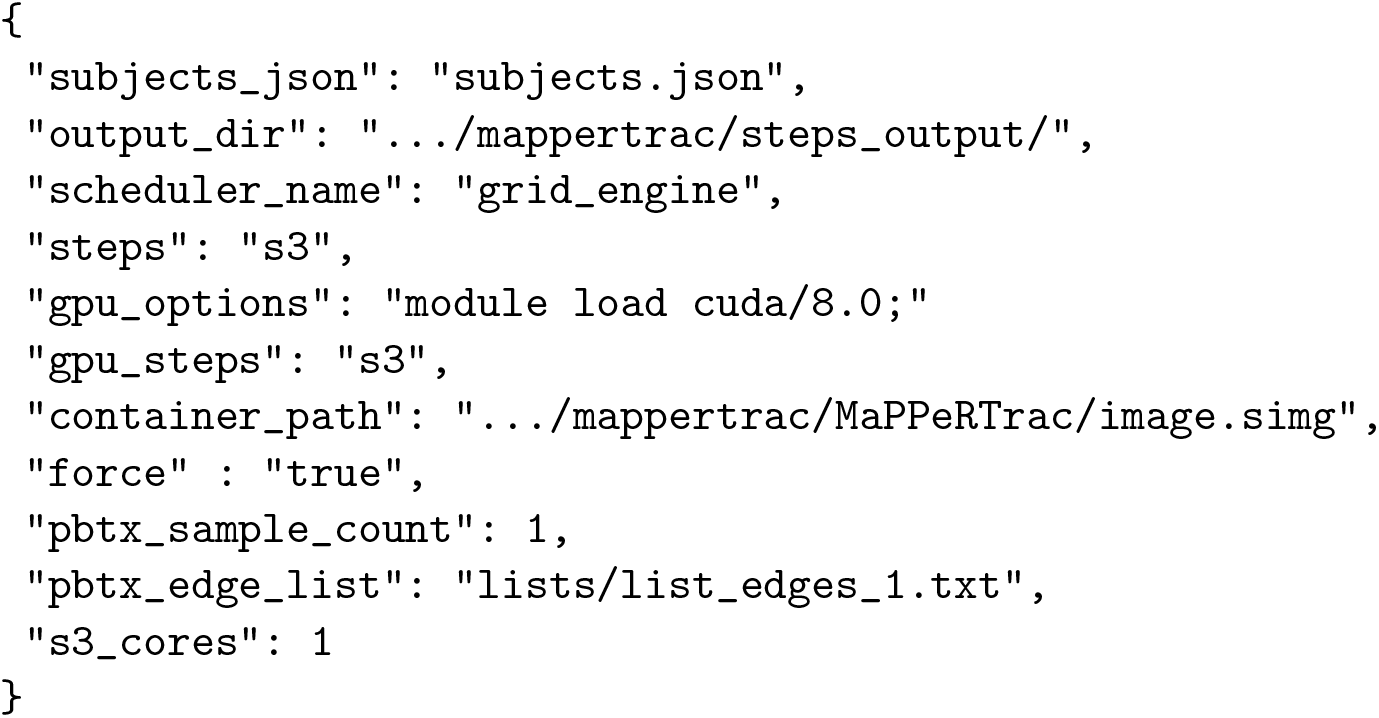

## Appendix D. Additional computational experiments

We experimented with many different approaches to speed-up file handling and tractography. The following experiments did not achieve sufficient speed-up for inclusion in the final software package.

1. *NVRAM disk to accelerate file IO* - we ran the pipeline using NVRAM memory on the Catalyst HPC cluster. This resulted in negligible performance improvement.
2. *Simplify tractography* - rather than run tractography between every single pair of regions, we attempted to run tractography a single time for each region and calculate connectivity post-hoc. However, this failed to capture sufficient edges and proved to be poorly supported by the PROBTRACKX2 software.
3. *Dynamic job times* - we tried to dynamically configure job times to help gain priority on computing queues. This proved unwieldy in practice.
4. *FreeSurfer GPU acceleration* - the slowest remaining part of the pipeline is FreeSurfer segmentation. We used pre-compiled binaries developed by (Hernandez-Fernandez *et al.*, 2019) to use GPU acceleration for FreeSurfer. However, this crashed on all Linux systems we tested in more than 90% of cases. Even when it ran successfully, speedup was only a modest 10-20%.

## Appendix E. Acknowledgements

The research is funded by the United States Department of Energy under the DOE Office of Science, Advanced Scientific Computing Research. Support is organized under The Co-Design for Artificial Intelligence and Computing at Scale for Extremely Large, Complex Datasets projects (Grant #KJ040301).

Geoffrey Manley and Pratik Mukherjee disclose grants from the United States Department of Defense – TBI Endpoints Development Initiative (Grant #W81XWH-14-2-0176), TRACK-TBI Precision Medicine (Grant #W81XWH-18-2-0042), and TRACK-TBI NETWORK (Grant #W81XWH-15-9-0001); NIH-NINDS – TRACK-TBI (Grant #U01NS086090); and the National Football League (NFL) Scientific Advisory Board – TRACK-TBI LONGITUDINAL.

The United States Department of Energy supports Dr. Manley for a precision medicine collaboration. One Mind has provided funding for TRACK-TBI patients stipends and support to clinical sites. He has received an unrestricted gift from the NFL to the UCSF Foundation to support research efforts of the TRACK-TBI NETWORK. Dr. Manley has also received funding from NeuroTruama Sciences LLC to support TRACK-TBI data curation efforts. Additionally, Abbott Laboratories has provided funding for add-in TRACK-TBI clinical studies.

Amy Markowitz receives funding from the Department of Defense TBI Endpoints Development Initiative (Grant #W81XWH-14-2-0176) and TRACK-TBI NETWORK (Grant #W81XWH-15-9-0001). Ms. Markowitz also receives salary support from the United States Department of Energy precision medicine collaboration and the philanthropic organization, One Mind.

## Footnotes

1 Transforming Research and Clinical Knowledge in Traumatic Brain Injury

2 https://hpc.llnl.gov/hardware/platforms/quartz

3 https://hpc.llnl.gov/hardware/platforms/pascal

4 https://hpc.llnl.gov/hardware/platforms/catalyst

5 https://wynton.ucsf.edu/hpc/about/specs.html

6 https://lcrc.anl.gov/systems/resources/blues

7 https://alcf.anl.gov/support-center/cooley/cooley-system-overview

8 https://aws.amazon.com/ec2/instance-types/p3

